# Network Architecture and Mutational Sensitivity of the *C. elegans* Metabolome

**DOI:** 10.1101/181511

**Authors:** Lindsay M. Johnson, Luke M. Chandler, Sarah K. Davies, Charles F. Baer

## Abstract

A fundamental issue in evolutionary systems biology is understanding the relationship between the topological architecture of a biological network, such as a metabolic network, and the evolution of the network. The rate at which an element in a metabolic network accumulates genetic variation via new mutations depends on both the size of the mutational target it presents and its robustness to mutational perturbation. Quantifying the relationship between topological properties of network elements and the mutability of those elements will facilitate understanding the variation in and evolution of networks at the level of populations and higher taxa.

We report an investigation into the relationship between two topological properties of 29 metabolites in the *C. elegans* metabolic network and the sensitivity of those metabolites to the cumulative effects of spontaneous mutation. The correlations between measures of network centrality and mutability are not statistically significant, but several trends point toward a weak positive association between network centrality and mutational sensitivity. There is a small but significant negative association between the mutational correlation of a pair of metabolites (*r*_*M*_) and the shortest path length between those metabolites.

Positive association between the centrality of a metabolite and its mutational heritability is consistent with centrally-positioned metabolites presenting a larger mutational target than peripheral ones, and is inconsistent with centrality conferring mutational robustness, at least *in toto*. The weakness of the correlation between *r*_*M*_ and the shortest path length between pairs of metabolites suggests that network locality is an important but not overwhelming factor governing mutational pleiotropy. These findings provide necessary background against which the effects of other evolutionary forces, most importantly natural selection, can be interpreted.

**Declarations:**

- <u>Ethics approval and consent to participate</u>: Not applicable
- <u>Consent for publication</u>: Not applicable
- <u>Availability of data and material</u>:
  - Metabolomics data (normalized metabolite concentrations) are archived in Dryad (http://dx.doi.org/10.5061/dryad.2dn09/1).
  - Data used to reconstruct the metabolic networks are included in Supplementary Appendix A1.
- <u>Competing interests</u>: The authors declare no competing interests.
- <u>Funding</u>: Funding was provided by NIH grant R01GM107227 to CFB and E. C. Andersen. The funding agency had no role in the design of the study and the collection, analysis, and interpretation of the data or in the writing of the manuscript.
- <u>Authors’ contributions</u>. LMJ and LMC collected and analyzed data in the network reconstruction and contributed to writing the manuscript. SKD collected and analyzed the GC-MS data. CFB analyzed data and wrote the manuscript. All authors read and approved the final manuscript.
- <u>Acknowledgements</u>: This work was initially conceived by Armand Leroi and Jake Bundy. We thank Art Edison, Dan Hahn, Tom Hladish, Marta Wayne, Michael Witting, and several anonymous reviewers for their generosity and helpful advice. We especially thank Hongwu Ma for leading us to and through his metabolite database and Reviewer #3 for his/her many insightful comments and suggestions. Support was provided by NIH grant R01GM107227 to CFB and E. C. Andersen.

## Introduction

The set of chemical reactions that constitute organismal metabolism is often represented as a network of interacting components, in which individual metabolites are the nodes in the network and the chemical reactions of metabolism are the edges linking the nodes (Jeong et al., 2000). Representation of a complex biological process such as metabolism as a network is conceptually powerful because it offers a convenient and familiar way of visualizing the system, as well as a well-developed mathematical framework for analysis.

If the representation of a biological system as a network is to be useful as more than a metaphor, it must have predictive power (Winterbach et al., 2013). Metabolic networks have been investigated in the context of evolution, toward a variety of ends. Many studies have compared empirical metabolic networks to various random networks, with the goal of inferring adaptive features of network architecture (e.g., Fell and Wagner, 2000;Jeong et al., 2000;Wagner and Fell, 2001;Siegal et al., 2007;Minnhagen and Bernhardsson, 2008;Papp et al., 2009;Bernhardsson and Minnhagen, 2010). Other studies have addressed the relationship between network-level properties of individual elements of the network (e.g., node degree, centrality) and properties such as rates of protein evolution (Vitkup et al., 2006;Greenberg et al., 2008), within-species polymorphism (Hudson and Conant, 2011), and mutational robustness (Levy and Siegal, 2008).

One fundamental evolutionary process that remains essentially unexplored with respect to metabolic networks is mutation. Mutation is the ultimate source of genetic variation, and as such provides the raw material for evolution: the greater the input of genetic variation by mutation, the greater the capacity for evolution. However, in a well-adapted population, most mutations are at least slightly deleterious. At equilibrium, the standing genetic variation in a population represents a balance between the input of new mutations that increase genetic variation and reduce fitness, and natural selection, which removes deleterious variants and thereby increases fitness. Because genetic variation is jointly governed by mutation and selection, understanding the evolution of any biological entity, such as a metabolic network, requires an independent accounting of the effects of mutation and selection.

The cumulative effects of spontaneous mutations can be assessed in the near absence of natural selection by means of a mutation accumulation (MA) experiment (Figure 1). Selection becomes ineffective relative to random genetic drift in small populations, and mutations with effects on fitness smaller than about the reciprocal of the population size (technically, the genetic effective population size, *N*_*e*_) will be essentially invisible to natural selection (Kimura, 1968). An MA experiment minimizes the efficacy of selection by minimizing *N*_*e*_, thereby allowing all but the most strongly deleterious mutations to evolve as if they are invisible to selection (Halligan and Keightley, 2009).

**Figure 1.**
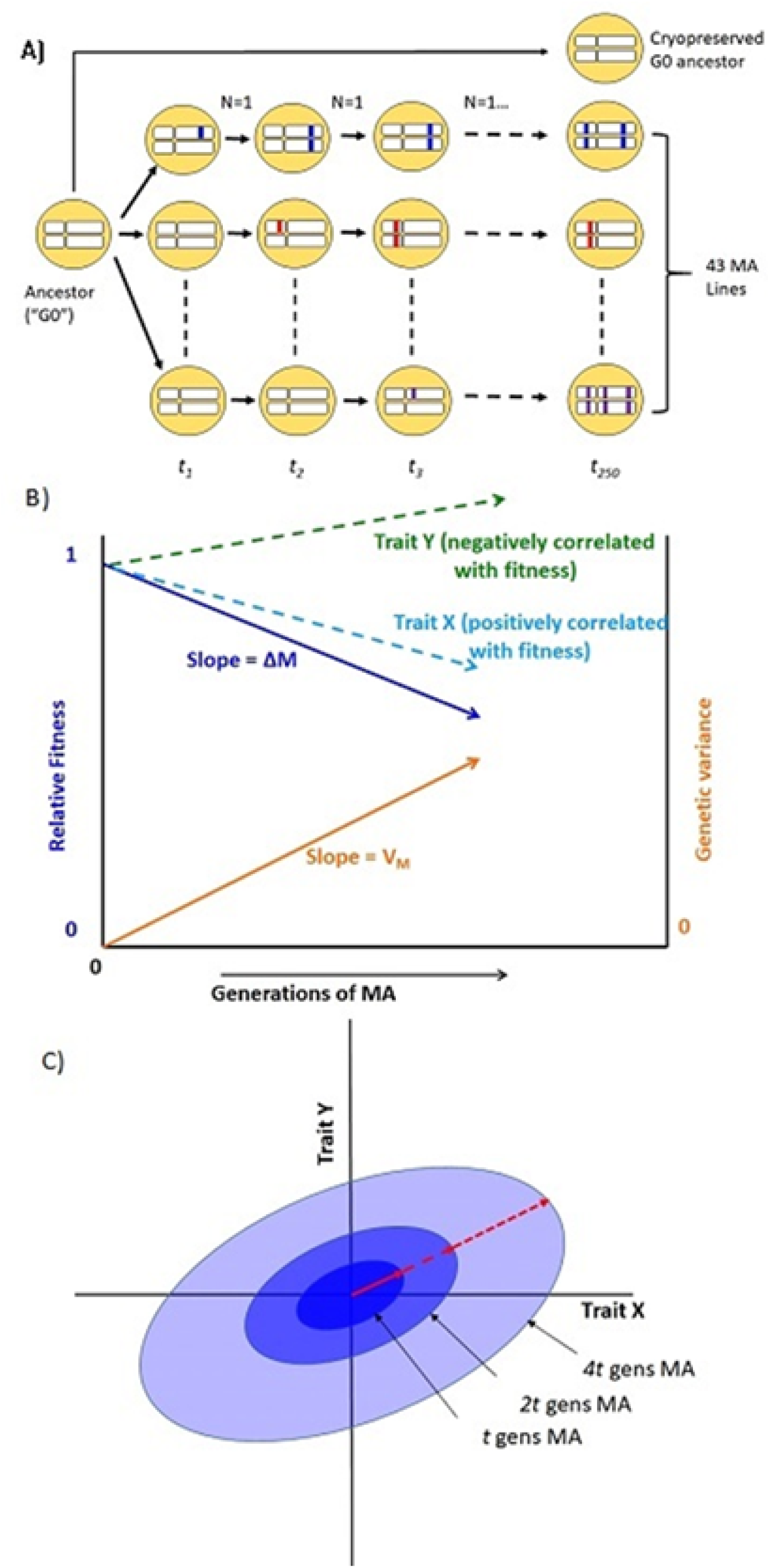
(**a**) Schematic diagram of the mutation accumulation (MA) experiment. An MA experiment is simply a pedigree. The genetically homogeneous ancestral line (G0) was subdivided into 100 MA lines, of which 43 are included in this study. Lines were allowed to accumulate mutations for *t*=250 generations. At each generation, lines were propagated by a single randomly chosen hermaphrodite (N=1). Mutations, represented as colored blocks within a homologous pair of chromosomes, arise initially as heterozygotes and are either lost or fixed over the course of the experiment. At the culmination of the experiment, each line has accumulated its own unique set of mutations. MA lines were compared to the cryopreserved G0 ancestor, which is wild-type at all loci. After Halligan and Keightley (2009). (**b**) Expected outcome of an MA experiment. As mutations accumulate over time, relative fitness (solid dark blue line) declines from its initial value of 1 at rate ΔM per generation and the genetic component of variance (solid orange line) increases from its initial value of 0 at rate V_M_ per generation. Trait X (light blue dashed line) is positively correlated with fitness and declines with MA; trait Y (green dashed line) is negatively correlated with fitness and increases with MA. Trajectories are depicted as linear, but they need not be. (**c**) Accumulation of mutational covariance in an MA experiment. Coordinate axes represent two traits, X and Y. Concentric ellipses show the increase in genetic covariance with MA, beginning from the initial value of zero; the orientation of the ellipses (red arrow) represents the linear relationship between pleiotropic mutational effects on the two traits.

Our primary interest is in the relationship between the centrality of a metabolite in the network and the sensitivity of that metabolite to mutation. Roughly speaking, the centrality of a node in a network quantifies some measure of the importance of the node in the network (Koschützki and Schreiber, 2008). A generic property of empirical networks, including metabolic networks, is that they are (approximately) scale-free; scale-free networks are characterized by a topology with a few “hub” nodes (high centrality) and many peripheral nodes (low centrality; Jeong et al., 2000). Scale-free networks are more robust to random perturbation than are randomly-connected networks (Albert et al., 2000).

Mutation is an important source of perturbation to biological systems, and much effort has gone into theoretical and empirical characterization of the conditions under which mutational robustness will evolve (Wagner et al., 1997;de Visser et al., 2003;Proulx et al., 2007). Mutational robustness can be assessed in two basic ways: top-down, in which a known element of the system is mutated and the downstream effects of the mutation quantified, or bottom-up, in which mutations are introduced at random, either spontaneously or by mutagenesis, and the downstream effects quantified. Top-down experiments are straightforward to interpret: the greater the effects of the mutation (e.g., on a phenotype of interest), the less robust the system. However, the scope of inference is limited to the types of mutations introduced by the investigator (which in practice are almost always gene knockouts), and provide limited insight into natural variation in mutational robustness.

Bottom-up approaches, in which mutations are allowed to accumulate at random, provide insight into the evolution of a system as it actually exists in nature: all else equal, a system, or element of a system (“trait”), that is robust to the effects of mutation will accumulate less genetic variance under MA conditions than one that is not robust (Figure 1b; Stearns et al., 1995). However, the inference is not straightforward, because all else may not be equal: different systems or traits may present different mutational targets (roughly speaking, the number of sites in the genome that potentially affect a trait; Houle (1998)).

Ultimately, disentangling the evolutionary relationship between network architecture, mutational robustness, and mutational target is an empirical enterprise, specific to the system of interest. As a first step, it is necessary to establish the relationship between network architecture (e.g., topology) and the rate of accumulation of genetic variance under MA conditions. If a general relationship emerges, targeted top-down experiments can then be employed to dissect the relationship in more mechanistic detail.

In addition to the relationship between metabolite centrality and mutational variance, we are also interested in the relationship between network topology and the mutational correlation (*r*_*M*_) between pairs of metabolites (Figure 1c). In principle, mutational correlations reflect pleiotropic relationships between genes underlying pairs of traits (but see below for caveats; Estes et al., 2005). Genetic networks are often modular (Newman, 2006), consisting of groups of genes (modules) within which pleiotropy is strong and between which pleiotropy is weak (Wagner et al., 2007). Genetic modularity implies that mutational correlations will be negatively correlated with the length of the shortest path between network elements. However, it is possible that the network of gene interactions underlying metabolic regulation is not tightly correlated with the metabolic network itself, e.g., if *trans* acting regulation predominates.

Here we report results from a long-term MA experiment in the nematode *Caenorhabditis elegans*, in which replicate MA lines derived from a genetically homogeneous common ancestor (G0) were allowed to evolve under minimally effective selection (*N*_*e*_≈1) for approximately 250 generations (Figure 1a). We previously reported estimates from these MA lines of two key quantitative genetic parameters by which the cumulative effects of mutation can be quantified: the per-generation change in the trait mean (the mutational bias, ΔM) and the per-generation increase in genetic variation (the mutational variance, V_M_) for the standing pools of 29 metabolites (Davies et al., 2016); Supplementary Table S1. In this report, we interpret those results, and new estimates of mutational correlations (*r*_*M*_), in the context of the topology of the *C. elegans* metabolic network.

## Methods and Materials

### I. Metabolic Network

The metabolic network of *C. elegans* was constructed following the criteria of Ma and Zeng (2003b), from two reaction databases *(i)* from Ma and Zeng (2003b); updated at http://www.ibiodesign.net/kneva/; we refer to this database as MZ, and *(ii)* from Yilmaz and Walhout (2016); http://wormflux.umassmed.edu/; we refer to this database as YW. Subnetworks that do not contain at least one of the 29 metabolites were excluded from downstream analyses. The method includes several *ad hoc* criteria for retaining or omitting specific metabolites from the analysis (criteria are listed on p. 272 of Ma and Zeng (2003b)). The set of reactions in the MZ and YW databases are approximately 99% congruent; in the few cases in which there is a discrepancy (listed in Supplementary Table S2), we chose to use the MZ database because we used the MZ criteria for categorizing currency metabolites (defined below).

To begin, the 29 metabolites of interest were identified and used as starting sites for the network. Next, all forward and reverse reactions stemming from the 29 metabolites were incorporated into the subnetwork until all reactions either looped back to the starting point or reached an endpoint. Currency metabolites were removed following the MZ criteria; a currency metabolite is roughly defined as a molecule such as water, proton, ATP, NADH, etc., that appears in a large fraction of metabolic reactions but is not itself an intermediate in an enzymatic pathway. Metabolic networks in which currency metabolites are included have much shorter paths than networks in which they are excluded. When currency metabolites are included in the network reported here, all shortest paths are reduced to no more than three steps, and most of the shortest paths consist of one or two steps. The biological relevance of path length when currency metabolites are included in the network is unclear (Ma and Zeng, 2003b).

A graphical representation of the network was constructed with the Pajek software package (http://mrvar.fdv.uni-lj.si/pajek/) and imported into the networkX Python package (Hagberg et al., 2008). Proper importation from Pajek to networkX was verified by visual inspection.

### II. Network Parameters

Properties of networks can be quantified in many ways, and different measures of network centrality capture different features of network importance (Table 1). We did not have strong prior hypotheses about which specific measure(s) of centrality associated with a given metabolite would prove most informative in terms of a relationship with the mutational properties of that metabolite (i.e., ΔM and/or V_M_). Therefore, we assessed the relationship between the mutational properties of a metabolite and several measures of its network centrality: betweenness, closeness, and degree centrality, in- and out-degree, and core number (depicted in Figure 3). These network parameters are all positively correlated. Definitions of the parameters are given in Table 1; correlations between the parameters are included in Table 2. Calculation of network parameters was done using built-in functions in NetworkX.

**Table 1.** Definitions of network parameters, following the documentation of NetworkX, v.1.11 (Hagberg et al. 2008) and mutational parameters. Abbreviations of the parameters used in Table 2 follow the parameter name in parentheses in bold type.

| Parameter | Heuristic Definition | Formal Definition |
| --- | --- | --- |
| In Degree (**IN**<sup>o</sup>), $deg^+(v)$ | The number of incoming edges to node $v$ in a directed graph. | self-explanatory |
| Out Degree (**OUT**<sup>o</sup>), $deg^-(v)$ | The number of outgoing edges from node $v$ in a directed graph. | self-explanatory |
| Shortest Path Length, $d(v, u)$ | Shortest distance from node $v$ to another node $u$ with no repeated walks | self-explanatory |
| Betweenness Centrality (**BET**), $c_B(v)$ | Betweenness centrality of node $v$ is the sum of the fraction of all-pairs shortest paths that pass through $v$. The greater $c_B(v)$, the greater the fraction of shortest paths that pass through node $v$. | $\frac{c_B(v)}{(n-1)(n-2)}$, where $c_B(v) = \sum_{s,t \in V} \frac{\sigma(s,t\vert v)}{\sigma(s,t)}$, $V$ is the set of nodes, $\sigma(s, t)$ is the number of shortest paths from node $s$ to node $t$, $\sigma(s, t\vert v)$ is the number of paths from $s$ to $t$ that pass through node $v$, and $n$ is the number of nodes in the graph. The denominator $(n-1)(n-2)$ is the normalization factor for a directed graph that scales $c_B(v)$ between 0 and 1. |
| Closeness Centrality (**CLO**),<br>$C(v)$ | Closeness centrality of node $v$ is the reciprocal of the sum of the shortest path lengths to all $n-1$ other nodes, normalized by the sum of minimum possible distances $n-1$. The greater $C(v)$, the closer $v$ is to other nodes. | $C(v) = \frac{n-1}{\sum_{u=1}^{n-1} d(u,v)}$, where $n$ is the number of nodes and $d(u, v)$ is the shortest path distance between $u$ and $v$. |
| Degree Centrality (**DEG**),<br>$C_D(v)$ | Degree centrality of node $v$ is the fraction of nodes in the network that node $v$ is connected to. | $C_D(v) = \frac{deg^+(v)+deg^-(v)}{n-1}$, where $n$ is the number of nodes in the network. |
| Core Number (**CORE**) | A $k$-core is the largest subgraph that contains nodes of at least degree $k$. The core number of node $v$ is the largest value $k$ of a $k$-core containing node $v$. | Calculated using the algorithm of Batagelj and Zaversnik (2011). |
| Mutational Bias (**ΔM**) | Per-generation rate of change of the trait mean in an MA experiment. Equivalent to the product of the genome-wide mutation rate, $\mu_G$, and the average effect of a mutation on the trait, $\alpha$. | $\Delta M_z = \frac{\bar{z}_{MA} - \bar{z}_0}{t\bar{z}_0}$; $\bar{z}_{MA}$ and $\bar{z}_0$ represent the MA and ancestral (G0) trait means and $t$ is the number of generations of MA. |
| Mutational Variance ($V_M$) | Per-generation rate of increase in genetic variance for a trait in an MA experiment. Equivalent to the product of the genome-wide mutation rate, $\mu_G$, and the square of the average effect of a mutation on the trait, $\alpha^2$. | $V_M = \Delta V_L = \frac{V_{L,MA} - V_{L,G0}}{2t}$, where $V_{L,MA}$ is the variance among MA lines, $V_{L,G0}$ is the among-line variance in the G0 ancestor, and $t$ is the number of generations of MA |
| Squared coefficient of variation ($I_M, I_E$) | $I_M$ is the mutational variance ($V_M$) scaled by the square of the trait mean, and provides a measure of the evolvability of a trait. $I_E$ is the residual variance ($V_E$) scaled in the same way. | |
| Mutational heritability ($h_M^2$) | Mutational variance ($V_M$) scaled as a fraction of the residual variance ($V_E$). Provides a measure of the short-term response to selection on mutational variance. | $h_M^2 = \frac{V_M}{V_E}$ |
| Mutational correlation (**$r_M$**) | Genetic correlation between two traits in MA lines. Provides an estimate of pleiotropic effects of new mutations. | $r_M = \frac{COV_M(X,Y)}{\sqrt{V_M(X)V_M(Y)}}$, where $COV_M$ is the mutational covariance and $V_M$ is the mutational variance. |

**Table 2.** Correlations between network parameters (Row/Column 1-5), between mutational parameters (Row/Column 6-9), between network and mutational parameters (shaded cells), and between residual variance (*I*_*E*_, Row/Column 10) and network and mutational parameters. Abbreviations of network parameters are: BTW, betweenness centrality; CLO, closeness centrality; DEG, degree centrality; IN°, in-degree, OUT°, out-degree; CORE, core number. Abbreviations of mutational parameters are: ΔM, per-generation change in the trait mean; |ΔM|, absolute value of ΔM; 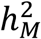, mutational heritability; *I*_*M*_, squared mutational CV; *I*_*E*_, squared residual CV. Network and mutational parameters are defined in Table 1. See text and Supplementary Table S1 for details of mutational parameters.* FDR < 0.1

|  | **BTW** | **CLO** | **DEG** | **IN°** | **OUT°** | **CORE** | **ΔM** | **\|ΔM\|** | **$h_M^2$** | **$I_M$** | **$I_E$** |
| --- | --- | --- | --- | --- | --- | --- | --- | --- | --- | --- | --- |
| **BTW** |  | 0.43 | 0.49 | 0.52 | 0.39 | 0.48 | -0.16 | -0.14 | 0.03 | -0.06 | -0.10 |
| **CLO** |  |  | 0.52 | 0.51 | 0.45 | 0.52 | 0.14 | 0.21 | 0.21 | 0.27 | 0.06 |
| **DEG** |  |  |  | 0.90 | 0.93 | 0.79 | 0.09 | 0.06 | 0.25 | 0.15 | 0.16 |
| **IN°** |  |  |  |  | 0.67 | 0.82 | 0.22 | 0.23 | 0.30 | 0.21 | 0.25 |
| **OUT°** |  |  |  |  |  | 0.64 | -0.04 | -0.08 | 0.17 | 0.09 | 0.05 |
| **CORE** |  |  |  |  |  |  | 0.33 | 0.28 | 0.53* | 0.30 | 0.28 |
| **ΔM** |  |  |  |  |  |  |  | 0.84 | 0.62 | 0.71 | 0.81 |
| **\|ΔM\|** |  |  |  |  |  |  |  |  | 0.53 | 0.69 | 0.84 |
| **$h_M^2$** |  |  |  |  |  |  |  |  |  | 0.72 | 0.43 |
| **$I_M$** |  |  |  |  |  |  |  |  |  |  | 0.82 |
| **$I_E$** |  |  |  |  |  |  |  |  |  |  |  |

### III. Mutation Accumulation Lines

A full description of the construction and propagation of the mutation accumulation (MA) lines is given in Baer et al. (2005). Briefly, 100 replicate MA lines were initiated from a nearly-isogenic population of N2-strain *C. elegans* and propagated by single-hermaphrodite descent at four-day (one generation) intervals for approximately 250 generations. The long-term *N*_*e*_ of the MA lines is very close to one, which means that mutations with a selective effect less than about 25% are effectively neutral (Keightley and Caballero, 1997). The common ancestor of the MA lines (“G0”) was cryopreserved at the outset of the experiment; MA lines were cryopreserved upon completion of the MA phase of the experiment. Based on extensive whole-genome sequencing (Denver et al., 2012; Saxena et al., submitted), we estimate that each MA line carries approximately 70 mutant alleles in the homozygous state.

At the time the metabolomics experiments reported in Davies et al. (2016) were initiated, approximately 70 of the 100 MA lines remained extant, of which 43 ultimately provided sufficient material for Gas Chromatography/Mass Spectrometry (GC-MS). Each MA line was initially replicated five-fold, although not all replicates provided data of sufficient quality to include in subsequent analyses; the mean number of replicates included per MA line is 3.9 (range = 2 to 5). The G0 ancestor was replicated nine times. However, the G0 ancestor was not subdivided into “pseudolines” (Teotónio et al., 2017), which means that inferences about mutational variances and covariances are necessarily predicated on the assumption that the among-line (co)variance of the ancestor is zero.

Each replicate consisted of stage-synchronized young adult worms taken from a single 10 cm agar plate. Cultures were stage-synchronized by treatment with hypochlorite (“bleaching”) following Stiernagle (2006); details of the synchronization are given in Davies et al. (2016). Following synchronization, worms were incubated at 20°C until young adulthood, defined as the point at which some eggs were seen on plates but no second generation worms had hatched. At this point, worms were washed from plates and collected for metabolomics. Each sample contained tens of thousands of worms, and although the samples were stage-synchronized, there was almost certainly some variation among samples in both the relative frequency of eggs on the plate and the (small) proportion of worms that had yet to reach adulthood.

Recently, whole-genome sequencing revealed that two MA lines, MA563 and MA564, share approximately 2/3 of their accumulated mutations; the simplest explanation is that the two lines were cross-contaminated around generation 150-175 of the MA protocol. However, averaged over all metabolites, the between-line standard deviation of those two lines is >3X that of either within-line SD, which suggests that the ∼1/3 of the mutations in each genome that are unique to each line contribute meaningfully to the differences between those two lines. Accordingly, we chose to include both lines. Further, since only 21 (out of 33) lines that we sequenced are represented in the metabolome dataset, the possibility of further unidentified cross-contamination cannot be ruled out. Comparisons between metabolites will not be biased by shared mutations, although the sampling (co)variance will increase by a factor 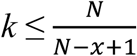 where *N* is the total number of lines and *x* is the number of lines that share mutations; 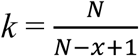 if all lines that share mutations share all their mutations.

### IV. Metabolomics

Details of the extraction and quantification of metabolites are given in Davies et al. (2016). Briefly, samples were analyzed using an Agilent 5975c quadrupole mass spectrometer with a 7890 gas chromatograph. Metabolites were identified by comparison of GC-MS features to the Fiehn Library (Kind et al., 2009) using the AMDIS deconvolution software (Halket et al., 1999), followed by reintegration of peaks using the GAVIN Matlab script (Behrends et al., 2011). Metabolites were quantified and normalized relative to an external quantitation standard. 34 metabolites were identified, of which 29 were ultimately included in the analyses. Normalized metabolite data are archived in Dryad (http://dx.doi.org/10.5061/dryad.2dn09).

### V. Mutational Parameters

In what follows, a “trait” is the (normalized) concentration of a metabolite. There are three mutational parameters of interest: *(i)* the per-generation proportional change in the trait mean, referred to as the mutational bias, ΔM; *(ii)* the per-generation increase in the genetic variance, referred to as the mutational variance, V_M_; and *(iii)* the genetic correlation between the cumulative effects of mutations affecting pairs of traits, the mutational correlation, *r*_*M*_. Details of the calculations of ΔM and V_M_ are reported in Davies et al. (2016); we reprise the basic calculations here.

*(i) Mutational bias (ΔM)* – The mutational bias is the change in the trait mean due to the cumulative effects of all mutations accrued over one generation. ΔM_z_=*µGα*_*z*_, where *µG* is the pergenome mutation rate and *α*_*z*_ is the average effect of a mutation on trait *z*, and is calculated as 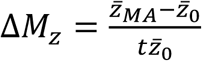, where 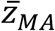 and 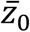 represent the MA and ancestral (G0) trait means and *t* is the number of generations of MA. However, the ΔM was not normally distributed among the 29 metabolites, so for downstream analyses we transformed ΔM as 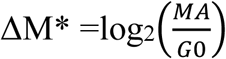, where MA and G0 represent the trait values of the MA lines and the G0 ancestor, respectively; ΔM=2^ΔM*^-1.

*(ii) Mutational variance (V*_*M*_) - The mutational variance is the increase in the genetic variance due to the cumulative effects of all mutations accrued over one generation.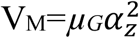 and is calculated as 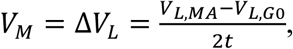, where *V*_*L,MA*_ is the variance among MA lines, *V*_*L,G0*_ is the among-line variance in the G0 ancestor, and *t* is the number of generations of MA (Lynch and Walsh, 1998, p. 330). In this study, we must assume that *V*_*L,G0*_ = 0.

Comparisons of variation among traits or groups require that the variance be measured on a common scale. V_M_ is commonly scaled either relative to the trait mean, in which case V_M_ is the squared coefficient of variation and is often designated *I*_*M*_, or relative to the residual variance, V_E_; V_M_/V_E_ is the mutational heritability, *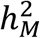*. *I*_*M*_ and 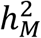 have different statistical properties and evolutionary interpretations (Houle et al., 1996), so we report both. For each metabolite, *I*_*M*_ and *I*_*E*_ are standardized relative to the mean of the MA lines. Both 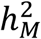 and *I*_*M*_ were natural-log transformed to meet assumptions of normality prior to downstream analyses.

*(iii) Mutational correlation, rM* – Pairwise mutational correlations were calculated from the among-line components of (co)variance, which were estimated by REML as implemented in the in the MIXED procedure of SAS v. 9.4, following Fry (2004). Statistical significance of individual correlations was assessed by Z-test, with a global 5% significance criterion of approximately P<0.000167.

### VI. Analysis of the relationship between mutational parameters and network centrality

The six network parameters are all positively correlated, as are the four mutational parameters (Table 2). To assess the overall correlation structure between mutational and network parameters, we employed a hierarchical canonical correlation analysis (CCA), as implemented in the CANCORR procedure of SAS v. 9.4, with the network parameters as the “X” variables and the mutational parameters as the “Y” variables. We initially included all four mutational parameters, resulting in four pairs of canonical variates and four canonical correlations. We then repeated the analysis for each mutational parameter *Y*_*i*_ individually with the full set of six network parameters, resulting in one pair of canonical variates and one canonical correlation for each of the four mutational parameters. Finally, we calculated the pairwise correlation between all mutational parameters and all network parameters. For all analyses except the first, significance was assessed using the False Discovery Rate (FDR) (Benjamini and Hochberg, 1995).

### IIV. Analysis of the relationship between mutational correlation (*r*_*M*_) and network architecture

*(i) Correlation between mutational correlation (rM) and shortest path length*. Statistical assessment of the correlation between mutational correlation (*r*_*M*_) and shortest path length presents a problem of non-independence, for two reasons. First, all correlations including the same variable (metabolite) are non-independent; each of the *n* elements of an *n* x *n* correlation matrix contributes to *n*(*n*-1)/2 correlations. Second, even though the mutational correlation between metabolites *i* and *j* is the same as the mutational correlation between *j* and *i*, the shortest path lengths need not be the same, and moreover, the path from *i* to *j* may exist whereas the path from *j* to *i* may not (depicted in Supplementary Figure S1). To account for non-independence of the data, we devised a parametric bootstrap procedure. Three metabolites (L-tryptophan, L-lysine, and Pantothenate) lie outside of the great strong component of the network (Ma and Zeng, 2003a) and are omitted from the analysis. Each off-diagonal element of the 24×24 mutational correlation matrix (*r*_*ij*_=*r*_*ji*_) was associated with a random shortest path length sampled with probability equal to its frequency in the empirical distribution of shortest path lengths between all metabolites included in the analysis. Next, we calculated the Spearman’s correlation *ρ* between *r*_*M*_ and the shortest path length. The procedure was repeated 10,000 times to generate an empirical distribution of *ρ*, to which the observed *ρ* can be compared. This comparison was done for the raw mutational correlation, *r*_*M*_, the absolute value, |*r*_*M*_|, and between *r*_*M*_ and the shortest path length in the undirected network (i.e., the shorter of the two paths between metabolites *i* and *j*).

## Results and Discussion

### Representation of the Metabolic Network

The metabolic network of *C. elegans* was estimated using method of Ma and Zeng (2003b) from two independent but largely congruent databases (Ma and Zeng, 2003b;Yilmaz and Walhout, 2016). Details of the network construction are given in section I of the Methods; data are presented in Supplementary Appendix A1. For the set of metabolites included (see Methods), networks constructed from the MZ and YW databases give nearly identical results. In the few cases in which there is a discrepancy (∼1%; Supplementary Table S2), we use the MZ network, for reasons we explain in the Methods. The resulting network is a directed graph including 646 metabolites, with 1203 reactions connecting nearly all metabolites (Figure 2).

**Figure 2.**
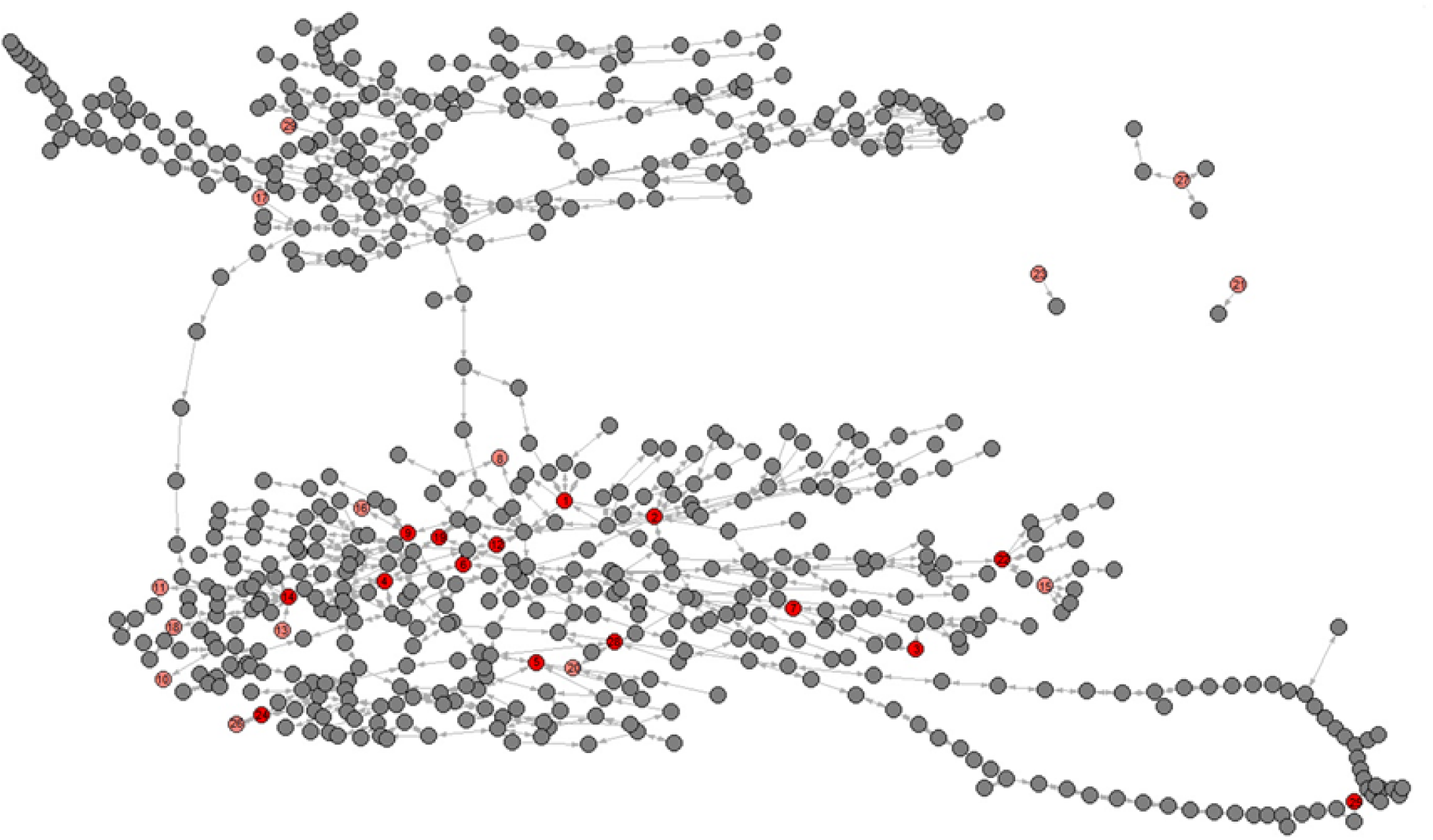
Graphical depiction of the metabolic network including all 29 metabolites. Pink nodes represent included metabolites with core number = 1, red nodes represent included metabolites with core number = 2. Gray nodes represent metabolites with which the included 29 metabolites directly interact. Metabolite identification numbers are: 1, L-Serine; 2, Glycine; 3, Nicotinate; 4, Succinate; 5, Uracil; 6, Fumarate; 7, L-Methionine; 8, L-Alanine. 9, L-Aspartate; 10, L-3-Amino-isobutanoate; 11, trans-4-Hydroxy-L-proline; 12, (S) – Malate; 13, 5-Oxoproline; 14, L-Glutamate; 15, L-Phenylalanine; `6, L-Asparagine; 17, D-Ribose; 18, Putrescine; 19, Citrate; 20, Adenine; 21, L-Lysine; 22, L-Tyrosine; 23, Pantothenate; 24, Xanthine; 25, Hexadecanoic acid; 26, Urate; 27, L-Tryptophan; 28, Adenosine; 29, Alpha;alpha-Trehalose.

**Figure 3.**
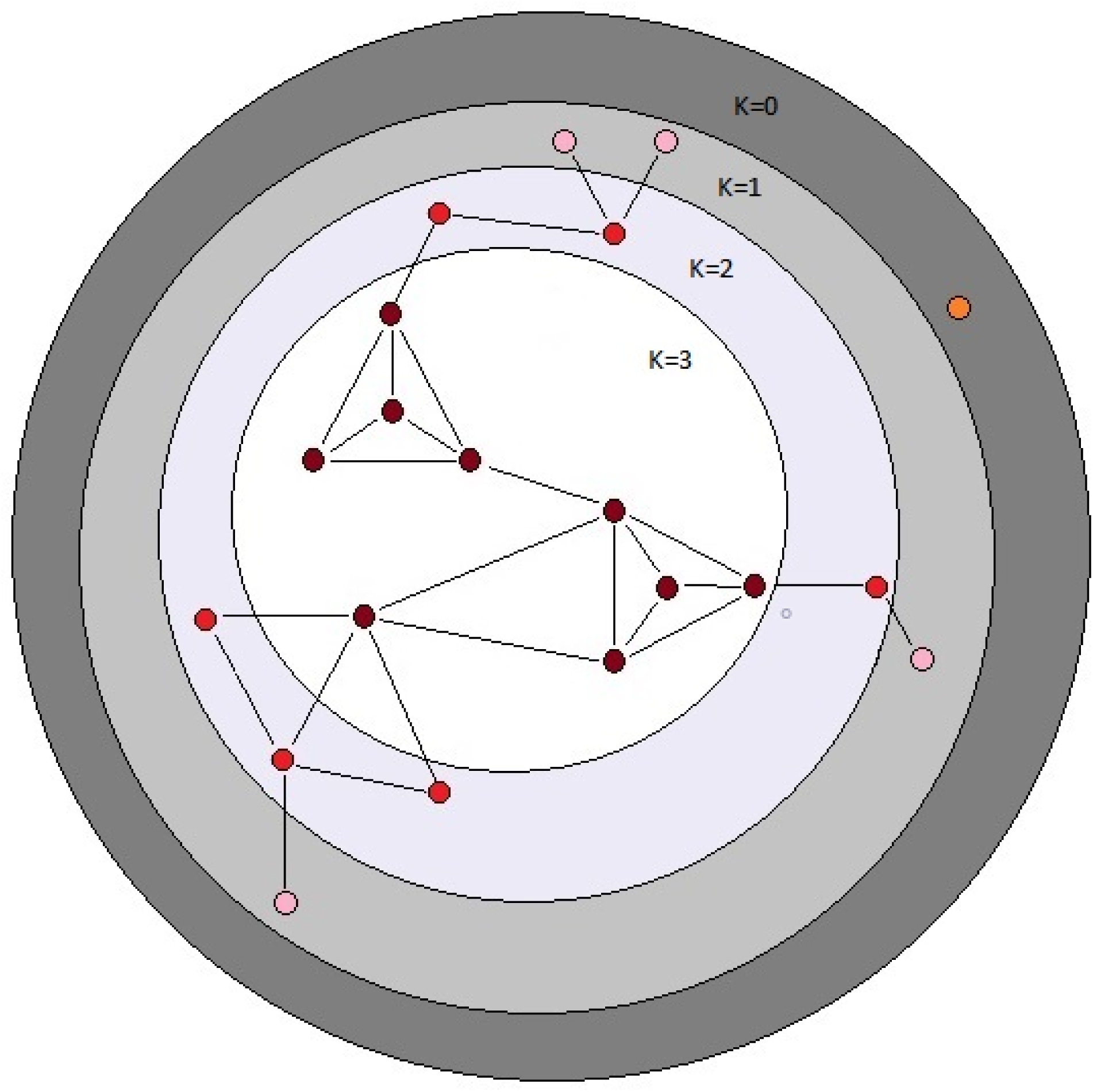
Schematic depiction of the *k*-cores of a graph. The *k*-core of a graph is the largest subgraph that contains nodes of degree at least *k*. The colored balls represent nodes in a network and the black lines represent connecting edges. Each dark red ball in the white area has core number *k*=3; note that each node with *k*=3 is connected to <u>at least</u> three other nodes. The depicted graph is undirected. After Batagelj and Zaversnik (2011).

### Network centrality and sensitivity to mutation

Canonical correlation analysis did not identify significant correlation between mutational parameters and network parameters, either collectively (Figure 4; Supplementary Table S3) or individually. Further, of the 24 pairwise correlations between mutational parameters and network parameters (Table 2, Supplementary Figure S2), only the correlation between mutational heritability (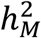) and core number approaches statistical significance (*r*=0.53, FDR < 0.1).

**Figure 4.**
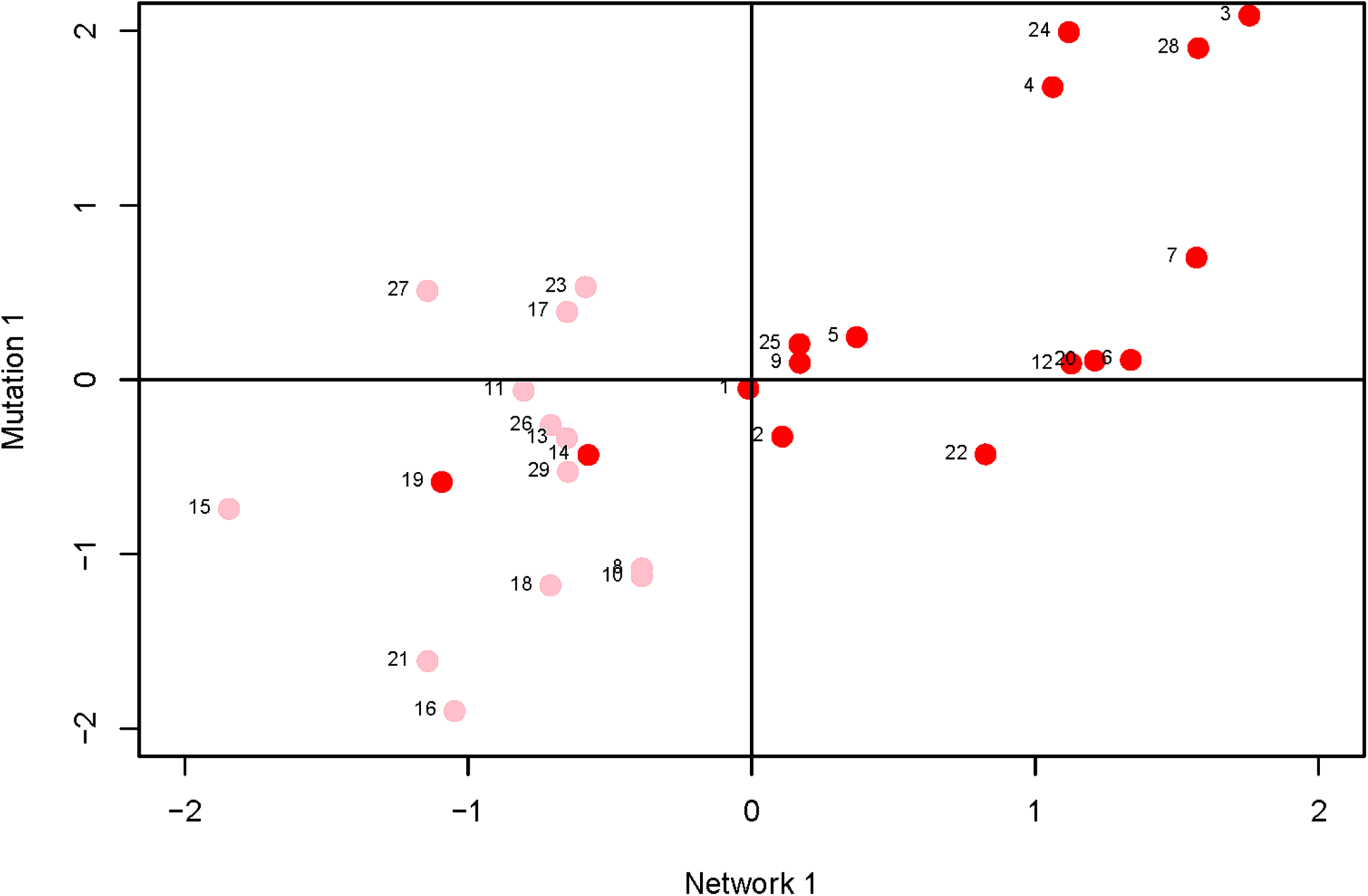
Plot of first canonical variate pair; the network variate is plotted on the X-axis, the mutation variate is plotted on the Y-axis. Each data point represents a metabolite; the numbers are the metabolite identifiers given in the legend to Figure 2. Metabolites with core number = 1 are in pink, metabolites with core number = 2 are in red.

On the face of it, it appears there is no association between network centrality and any measure of mutational sensitivity. If so, there are various possible explanations. For example, it may be that mutational target and mutational robustness effectively cancel each other out. More worryingly, it may be that the representation of the *C. elegans* metabolic network used here misrepresents the network as it actually exists *in vivo*. For example, the topology of the dynamic metabolic network of the bacterium *E. coli* varies depending on the environmental context (Koschützki et al., 2010), and it seems intuitive that the greater spatiotemporal complexity inherent to a multicellular organism would exacerbate that problem. Or, most straightforwardly, it may be that there simply is no functional relationship between the centrality of a metabolite in a network and its sensitivity to mutation.

However, several trends apparent in the results suggest the conservative interpretation may miss meaningful signal emerging from noisy data. First, the point estimates of the canonical correlations are not small (> 0.45 in all five cases; e.g., the first canonical correlation in the full analysis is 0.69; Supplementary Table 3); it may simply be that the sampling variance associated with the relatively small number of mutations, MA lines and (especially) metabolites overwhelms the signal of a weak but consistently positive association. Second, of the 24 pairwise correlations among mutational and network parameters (Table 2), only five are negative, significantly fewer than expected at random if the variables are uncorrelated (cumulative binomial probability = 0.0033). Third, the point estimates of the pairwise correlations are not random with respect to either network or mutational parameters. For all four mutational parameters, the correlation is greatest with core number (exact probability ≈ 0.00077). Core number is a discrete interval variable, whereas the other measures of network centrality are continuous variables. Quantifying centrality in terms of core number is analogous to categorizing a set of size measurements into “small” and “large”: power is increased, at the cost of losing the ability to discriminate between subtler differences.

Fourth, for five out of six network parameters, the correlation is greatest with 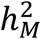 (exact cumulative probability ≈ 0.00066). V_M_ is the numerator of both 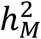 and *I*_*M*_; the difference is the denominator, with 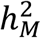 scaling V_M_ by the residual variance, V_E_, and *I*_*M*_ scaling V_M_ by the square of the trait mean. If V_E_ was more strongly associated with network topology than was V_M_, 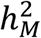 would presumably be more strongly correlated with network parameters than would *I*_*M*_, analogous to the well-documented V_E_-driven negative association between the narrow-sense heritability of a trait and the correlation of the trait with fitness (Houle, 1992). However, *I*_*M*_ and *I*_*E*_ are nearly identically (un)correlated with network parameters (Table 2), so that scenario cannot explain the correlation. Coincidence seems as likely an explanation as any.

### The relationship between mutational correlation (*r*_*M*_) and shortest path length

In an MA experiment, the cumulative effects of mutations on a pair of traits *i* and *j* may covary for two, nonexclusive reasons (Estes et al., 2005). More interestingly, individual mutations may have consistently pleiotropic effects, such that mutations that affect trait *i* also affect trait *j* in a consistent way. Less interestingly, but unavoidably, individual MA lines will have accumulated different numbers of mutations, and if mutations have consistently directional effects, as would be expected for traits correlated with fitness, lines with more mutations will have more extreme trait values than lines with fewer mutations, even in the absence of consistent pleiotropy. Estes et al. (2005) simulated the sampling process in *C. elegans* MA lines with mutational properties derived from empirical estimates from a variety of traits and concluded that sampling is not likely to lead to large absolute mutational correlations in the absence of consistent pleiotropy (|*r*_*M*_| ≤ 0.25).

Ideally, we would like to estimate the full mutational (co)variance matrix, ***M***, from the joint estimate of the among-line (co)variance matrix. However, with 25 traits, there are (25×26)/2 = 325 covariances, and with only 43 MA lines, there is insufficient information to jointly estimate the restricted maximum likelihood of the full ***M*** matrix. To proceed, we calculated mutational correlations from pairwise REML estimates of the among-line (co)variances, i.e., 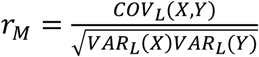 (Clark et al., 1995;Mezey and Houle, 2005). Pairwise estimates of *r*_*M*_ are shown in Supplementary Table S4. To assess the extent to which the pairwise correlations are sensitive to the underlying covariance structure, we devised a heuristic bootstrap analysis. For a random subset of 12 of the 300 pairs of traits, we randomly sampled six of the remaining 23 traits without replacement and estimated *r*_*M*_ between the two focal traits from the joint REML among-line (co)variance matrix. For each of the 12 pairs of focal traits, we repeated the analysis 100 times.

There is a technical caveat to the preceding bootstrap analysis. Resampling statistics are predicated on the assumption that the variables are exchangeable (Shaw, 1992), which metabolites are not. For that reason, we do not present confidence intervals on the resampled correlations, only the distributions. However, we believe that the analysis provides a meaningful heuristic by which the sensitivity of the pairwise correlations to the underlying covariance structure can be assessed.

Distributions of resampled correlations are shown in Supplementary Figure S3. In every case the point estimate of *r*_*M*_ falls on the mode of the distribution of resampled correlations, and in 11 of the 12 cases, the median of the resampled distribution is very close to the point estimate of *r*_*M*_. However, in six of the 12 cases, some fraction of the resampled distribution falls outside two standard errors of the point estimate. The most important point that the resampling analysis reveals is this: given that 29 metabolites encompass only a small fraction of the total metabolome of *C. elegans* (<5%), even had we been able to estimate the joint likelihood of the full 29×30/2 ***M***-matrix, the true covariance relationships among those 29 metabolites could conceivably be quite different from those estimated from the data.

The simplest property that describes the relationship between two nodes in a network is the length of the shortest path between them (= number of edges). In a directed network, such as a metabolic network, the shortest path from element *i* to element *j* is not necessarily the same as the shortest path from *j* to *i*. For each pair of metabolites *i* and *j*, we calculated the shortest path length from *i* to *j* and from *j* to *i*, without repeated walks (Supplementary Table S5). We then calculated Spearman’s correlation *ρ* between the mutational correlation *r*_*M*_ and the shortest path length.

There is a weak, but significant, negative correlation between *r*_*M*_ and the shortest path length between the two metabolites (*ρ* = −0.128, two-tailed P<0.03; Figure 5a), whereas |*r*_*M*_| is not significantly correlated with shortest path length (*ρ* = −0.0058, two-tailed P>0.45; Supplementary Figure 5b). The correlation between *r*_*M*_ and the shortest path in the undirected network is similar to the correlation between *r*_*M*_ and the shortest path in the directed network (*ρ* = −0.105, two-tailed P>0.10; Supplementary Figure 5c).

**Figure 5.**
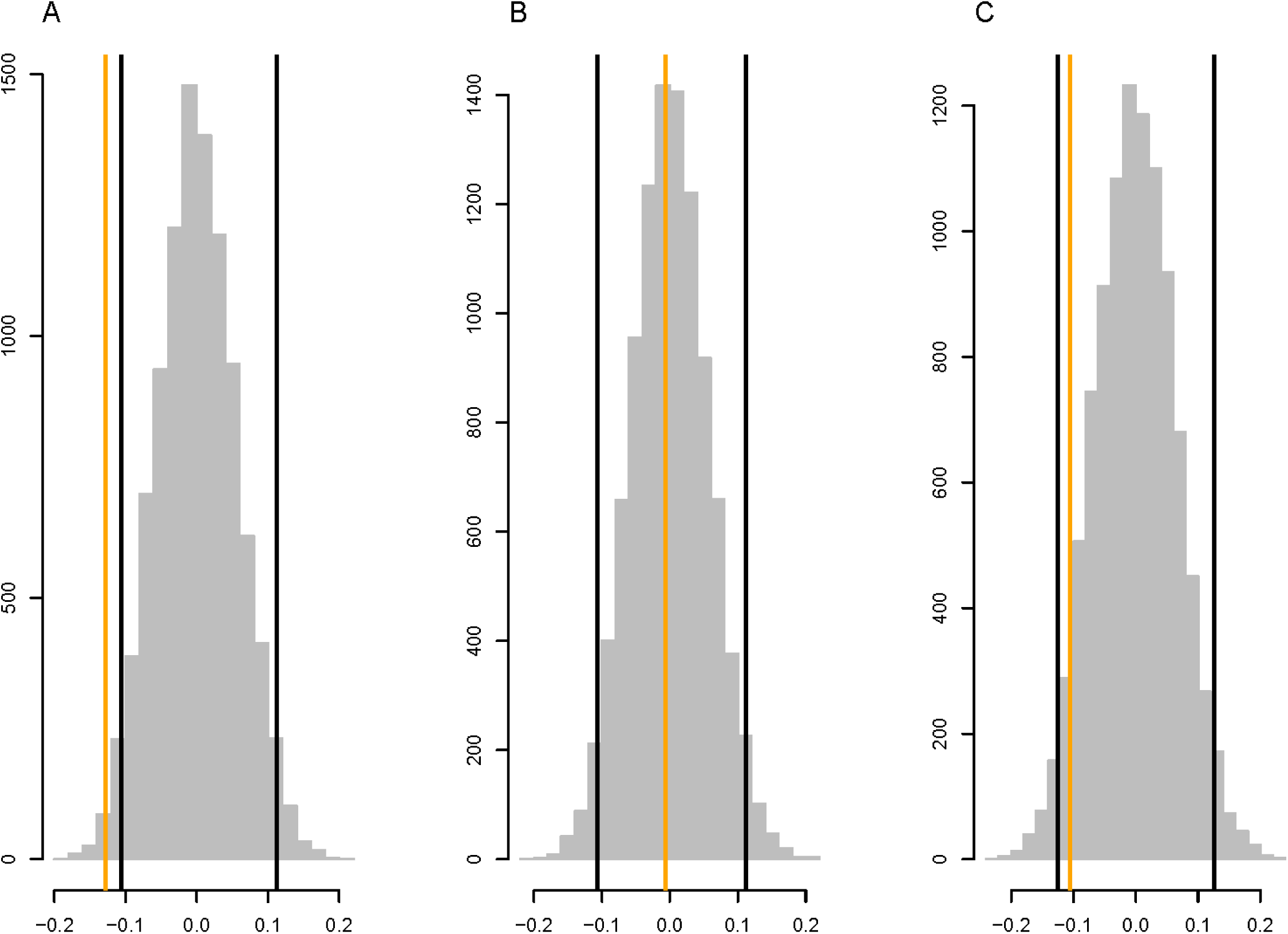
Parametric bootstrap distributions of random correlations *ρ* between (**a**) *r*_*M*_ and the shortest path length in the directed network, (**b**) |*r*_*M*_| and the shortest path length in the directed network, (**c**) *r*_*M*_ and shortest path length in the undirected network (i.e., the shorter of the two path lengths between metabolites *i* and *j* in the directed network). Orange lines show the observed values of *ρ*, black lines show the 95% confidence interval of the distribution of the correlation between the mutational correlation and a random shortest path length drawn from the observed distribution of shortest path lengths. See Methods for details.

An intuitive possible cause of the weak negative association between shortest path length and mutational correlation would be if a mutation that perturbs a metabolic pathway toward the beginning of the pathway has effects that propagate downstream in the same pathway, but the effect of the perturbation attenuates. The attenuation could be due either to random noise or to the effects of other inputs into the pathway downstream from the perturbation (or both). The net effect would be a characteristic pathway length past which the mutational effects on two metabolites are uncorrelated, leading to an overall negative correlation between *r*_*M*_ and path length. The finding that the correlations between *r*_*M*_ and the shortest path length in the directed and undirected network are very similar reinforces that conclusion. The negative correlation between *r*_*M*_ and shortest path length is reminiscent of a finding from Arabidopsis, in which sets of metabolites significantly altered by single random gene knockouts are closer in the global metabolic network than expected by chance (Kim et al., 2015).

### Conclusions and Future Directions

The proximate goal of this study was to find out if there are topological properties of the *C. elegans* metabolic network (node centrality, shortest path length) that are correlated with a set of statistical descriptions of the cumulative effects of spontaneous mutations (ΔM, V_M_, *r*_*M*_). Ultimately, we hope that a deeper understanding of those mathematical relationships will shed light on the mechanistic biology of the organism. Bearing in mind the statistical fragility of the results, we conclude:

*(i) Network centrality <u>may</u> be associated with mutational sensitivity (V*_*M*_*); it is not associated with mutational robustness (1/V*_*M*_). If in fact the apparently non-random features of the data represent a hint of signal emerging from the noise, the most plausible explanation is that metabolites that are central in the network present a larger mutational target than do metabolites that peripherally located. Somewhat analogously, Landry et al. (2007) investigated the mutational properties of transcription in a set of yeast MA lines and found that 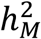 is positively correlated with both the number of genes with which a given gene interacts (“trans-mutational target”) and the number of transcription factor binding sites in a gene’s promoter (“cis-mutational target”). Those authors did not formally quantify the network properties of the set of transcripts, although is seems likely that mutational target size as they defined it is positively correlated with centrality in the transcriptional network. It is important to note, however, although 1/V_M_ is a meaningful measure of mutational robustness (Stearns and Kawecki, 1994), it does not necessarily follow that highly-connected metabolites are therefore more robust to the effects of *individual* mutations (Houle, 1998;Ho and Zhang, 2016).

*(ii) Pleiotropic effects of mutations affecting the metabolome are predominantly local*, as evidenced by the significant negative correlation between the mutational correlation, *r*_*M*_, and the shortest path length between a pair of metabolites. That result is not surprising in hindsight, but the weakness of the correlation suggests that there are other important factors that underlie pleiotropy beyond network proximity.

*(iii) Future Directions*. To advance understanding of the mutability of the *C. elegans* metabolic network, three things are needed. First, it will be important to cover a larger fraction of the metabolic network. Untargeted mass spectrometry of cultures of *C. elegans* reveals many thousands of features (Art Edison, personal communication); 29 metabolites are only the tip of a large iceberg. For example, our intuition leads us to believe that the mutability of a metabolite will depend more on its in-degree (mathematically, the number of edges leading into a node in a directed graph; biochemically, the number of reactions in which the metabolite is a product) than its out-degree. For all four mutational parameters, the point-estimate of the pairwise correlation with in-degree is greater than that with out-degree (Table 2), although that result is not statistically significant (binomial probability = 0.0625).

Second, to more precisely partition mutational (co)variance into within- and among-line components, more MA lines are needed. We estimate that each MA line carries about 70 unique mutations (see Methods), thus the mutational (co)variance is the result of about 3000 total mutations, distributed among 43 MA lines. The MA lines were a preexisting resource, and the sample size was predetermined. It is encouraging that we were able to detect significant mutational variance for 25/29 metabolites (Supplementary Table S1), but only 14% (42/300) of pairwise mutational correlations are significantly different from zero at the experiment-wide 5% significance level, roughly corresponding to |*r*_*M*_|>0.5 (Supplementary Table S4); 18 of the 42 significant mutational correlations are not significantly different from |*r*_*M*_| = 1. It remains uncertain how sensitive estimates of mutational correlations are to the underlying covariance structure of the metabolome. It also remains to be seen if the mutability of specific features of metabolic networks are genotype or species-specific, and the extent to which mutability depends on environmental context.

Third, it will be important to quantify metabolites (static concentrations and fluxes) with more precision. The metabolite data analyzed in this study were collected from large cultures (n>10,000 individuals) of approximately stage-synchronized worms, and were normalized relative to an external quantitation standard (Davies et al., 2016). Ideally, one would like to characterize the metabolomes of single individuals, assayed at the identical stage of development. Single-worm metabolomics is on the near horizon (M. Witting, personal communication). Minimizing the number of individuals in a sample is important for two reasons; (1) the smaller the sample, the easier it is to be certain the individuals are at the same developmental stage, and (2) knowing the exact number of individuals in a sample makes normalization relative to an external standard more interpretable. Ideally, data would be normalized relative to both an external standard and an internal standard (e.g., total protein; Clark et al. (1995)).

This study provides an initial assessment of the relationship between mutation and metabolic network architecture. To begin to uncover the relationship between metabolic architecture and natural selection, the next step is to repeat these analyses with respect to the standing genetic variation (V_G_). There is some reason to think that more centrally-positioned metabolites will be more evolutionarily constrained (i.e., under stronger purifying selection) than peripheral metabolites (Vitkup et al., 2006), in which case the ratio of the mutational variance to the standing genetic variance (V_M_/V_G_) will increase with increasing centrality.

## List of Figures and Tables

**Supplementary Figure S1.** Depiction of shortest path length in a directed network.

**Supplementary Figure S2.** Plots of mutational parameters vs. network statistics.

**Supplementary Figure S3.** Bootstrap distributions of *rM* with six randomly chosen covariates.

**Supplementary Table S1.** Network and mutational parameters of metabolites. (Excel)

**Supplementary Table S2.** Table of discrepancies between MZ and YW methods (Word)

**Supplementary Table S3.** Canonical correlation analysis (Word).

**Supplementary Table S4.** Mutational and environmental correlations (Excel)

**Supplementary Table S5.** Shortest network path lengths (Excel)

**Supplementary Data Set 1.** Metabolic network data (Excel)

## Acknowledgments

This work was initially conceived by Armand Leroi and Jake Bundy. We thank Art Edison, Dan Hahn, Tom Hladish, Marta Wayne, Michael Witting, and several anonymous reviewers for their generosity and helpful advice. We especially thank Hongwu Ma for leading us to and through his metabolite database and Reviewer #3 for his/her many insightful comments and suggestions. Support was provided by NIH grant R01GM107227 to CFB and E. C. Andersen.

