## Supplementary Materials for "Network Architecture and Mutational Sensitivity of the *C. elegans* Metabolome" (from another version of this preprint: version 4, 2018-03-25)

| **Reaction Number** | **Reaction** |
| --- | --- |
| R01579 | D-Glutamine + H2O = D-Glutamate + NH3 |
| R01887 | gamma-Amino-gamma-cyanobutanoate + 2 H2O = DL-Glutamate + NH3 |
| R04936 | Se-Adenosylselenohomocysteine + H2O = Adenosine +Selenohomocysteine |
| R00891 | L-Serine + Hydrogen sulfide = L-Cysteine + H2O |
| R09099 | L-Serine + 5,6,7,8-Tetrahydromethanopterin = 5,10-Methylenetetrahydromethanopterin + Glycine + H2O |
| R02853 | D-O-Phosphoserine + H2O = D-Serine + Orthophosphate |
| R00904 | 3-Aminopropanal + NAD+ + H2O = beta-Alanine + NADH + H+ |
| R03542 | alpha-Aminopropiononitrile + 2 H2O = Alanine + NH3 |
| R01324 | Citrate = Isocitrate |
| R00483 | ATP + L-Aspartate + NH3 = AMP + Diphosphate + L-Asparagine |
| R01221 | Glycine + Tetrahydrofolate + NAD+ = 5,10-Methylenetetrahydrofolate+ NH3 + CO2 + NADH + H+ |
| R02078 | 3,4-Dihydroxy-L-phenylalanine + L-Tyrosine + Oxygen = Dopaquinone+ 3,4-Dihydroxy-L-phenylalanine + H2O |
| R01706 | Hexadecanoyl-[acp] + H2O = Acyl-carrier protein + Hexadecanoicacid |
| R04666 | 3-Ureidoisobutyrate + H2O = 3-Aminoisobutyric acid + CO2 + NH3 |

**Supplementary Table S2.** Discrepancies between the metabolic networks constructed using the MZ and YW reaction databases. All reactions listed here are in the Ma and Zeng database (<http://www.ibiodesign.net/kneva/>) but not in the Wormflux database (<http://wormflux.umassmed.edu/>) and were used in the generation of the metabolic network. There is a total of 1203 reactions in the network, these represent about 1% of all reactions.
